# scDeepInsight: a supervised cell-type identification method for scRNA-seq data with deep learning

**DOI:** 10.1101/2023.03.09.531861

**Authors:** Shangru Jia, Artem Lysenko, Keith A Boroevich, Alok Sharma, Tatsuhiko Tsunoda

**Affiliations:** Laboratory for Medical Science Mathematics, Department of Computational Biology and Medical Sciences, Graduate School of Frontiwer Sciences, The University of Tokyo, Japan; Laboratory for Medical Science Mathematics, Department of Biological Sciences, School of Science, The University of Tokyo, Japan; Laboratory for Medical Science Mathematics, RIKEN Center for Integrative Medical Sciences, Japan; Institute for Integrated and Intelligent Systems, Griffith University, Australia

**Keywords:** single-cell RNA sequencing, deep learning, cell annotation, transformers

## Abstract

Annotation of cell-types is a critical step in the analysis of single-cell RNA sequencing (scRNA-seq) data that allows the study of heterogeneity across multiple cell populations. Currently this is most commonly done using unsupervised clustering algorithms, which project single-cell expression data into a lower dimensional space and then cluster cells based on their distances from each other. However, as these methods do not use reference datasets, they can only achieve a rough classification of cell-types, and it is difficult to improve the recognition accuracy further. To effectively solve this issue we propose a novel supervised annotation method, scDeepInsight. The scDeepInsight method is capable of performing manifold assignments. It is competent in executing data integration through batch normalization, performing supervised training on the reference dataset, doing outlier detection and annotating cell-types on query datasets. Moreover, it can help identify active genes or marker genes related to cell-types. The training of the scDeepInsight model is performed in a unique way. Tabular scRNA-seq data are first converted to corresponding images through the DeepInsight methodology. DeepInsight can create a trainable image transformer to convert non-image RNA data to images by comprehensively comparing interrelationships among multiple genes. Subsequently, the converted images are fed into convolutional neural networks (CNNs) such as EfficientNet-b3. This enables automatic feature extraction to identify the cell-types of scRNA-seq samples. We benchmarked scDeepInsight with six other mainstream cell annotation methods. The average accuracy rate of scDeepInsight reached 87.5%, which is more than 7% higher compared with the state-of-the-art methods.

## Introduction

Rapidly developing single-cell RNA sequencing (scRNA-seq) technologies have made it possible to observe gene expression at the single-cell level and enhance our understanding of complex biological systems and diseases, such as cancer and chronic diseases. These methods have significant implications for studying a wide variety of tissues and the different types of cells within them. Accurate cell annotation is a prerequisite for downstream analysis of single-cell data. However, cell annotation is a time-consuming and expert-dependent step in single-cell analysis. The annotation process can be divided into three steps [1], automatic annotation, manual annotation by an expert and verification. Due to the emergence of a large number of single-cell datasets, manual annotation by experts based on empirical analysis is practically impossible at a rate needed to meet the research needs. Furthermore, manual annotation is not only time-consuming and laborious, but the annotation results can be subjective.

Owing to these factors, improving the accuracy of automatic annotation has become an essential area of research and in response to these practical needs the researchers have proposed a variety of relevant annotation methods. These can be roughly divided into three different types according to the theoretical basis [2]: 1) annotation based on marker genes, 2) annotation using correlation analysis with reference datasets, and 3) annotation by supervised classifiers trained on reference datasets.

Most single-cell annotation methods start with unsupervised cell clustering analysis. Cells are first assigned into groups using clustering methods such as k-means [3], Single-Cell Consensus Clustering (SC3) [4], and shared nearest neighbor (SNN) [5]. The clusters are then mapped to different cell-types by analyzing the abundance of marker genes within each cluster. However, these marker gene-based methods have several limitations. The first issue is the accuracy of the marker gene database. Even though there are now databases such as PanglaoDB [6], ScType [7], and CellMarker [8], the selection of some marker genes still depends on prior research knowledge. The second issue is that information on marker genes is often insufficient for many cell subtypes and, in particular, newly discovered cell-types. The third issue is the duplication of marker genes between subtypes of cells. Taking the ScType database as an example, the marker genes of multiple subclasses of B cells heavily overlap with each other, and the lack of specificity leads to confusion between similar subtypes during classification.

Annotation methods based on correlation analysis with reference datasets tend to be more accurate than marker gene-based methods [9]. This is because gene-gene correlations are generally ignored when analyzing marker gene lists. Annotation can be done more comprehensively by correlating target unannotated datasets with reference datasets of similar biological tissues. However, batch effects between the reference dataset and the target dataset can hinder the correct annotation of cell-types. Technical differences, such as sequencing methods and experimental batches, will affect the results of single-cell sequencing. Despite this, current correlation-based annotation methods, such as SingleR [10], often do not offer batch effect processing methods as part of their pipeline. When annotating datasets obtained in different experiments, it is very difficult to eliminate the influence of the batch effects.

Single-cell datasets are often high-dimensional and sparse. In this case, machine learning (ML) is a good choice for processing complex sequencing data. After learning the expression patterns of multiple genes in different cell-types on the reference dataset, ML methods can transfer labels from the reference dataset to the target dataset. Deep learning methods in particular are capable of learning highly abstracted representations from data such as images, sounds, and texts. Given the robustness of deep learning methods and the availability of finely annotated reference datasets, supervised learning models have gradually become widely used in the analysis of reference datasets. Currently, representative machine learning methods for processing single-cell omics data mainly include Bidirectional Encoder Representations from Transformers (BERT) [2], Autoencoders (AEs) [11] and Recurrent Neural Networks (RNNs) [12]. Single-cell omics data are often treated as texts or sequence data since they are not images and do not have graph network structures.

In this paper, we propose scDeepInsight, an original method integrating the whole cell-type identification process (Figure 1). scDeepInsight can directly annotate the query dataset based on the model trained on the reference dataset. In the first step, scDeepInsight does preprocessing of scRNA-seq data, including quality control and integration through batch normalization. By incorporating DeepInsight [13], our method can convert the scRNA-seq samples into corresponding images. Images generated from the reference dataset are used to train a CNN, which can then be used to predict cell types found in the query dataset(s). The absolute superiority of CNNs in image classification and feature extraction has been widely recognized [14]. Furthermore, extracted features in the training process are helpful in investigating marker genes by using DeepFeature [15]. With a unique approach for converting scRNA-seq to image data, our method allows to fully exploit the advantages of CNNs in cell-type classification. scDeepInsight enables accurate and efficient annotation of multiple cell subtypes and can perform outlier detection. Cell-type prediction results are validated using reliable pre-annotated cell labels. Also, some rare/unknown cell-types, which are not included in the reference datasets, can be detected during the annotation process. Further details are explained in the Materials and Methods section.

**Figure 1.**
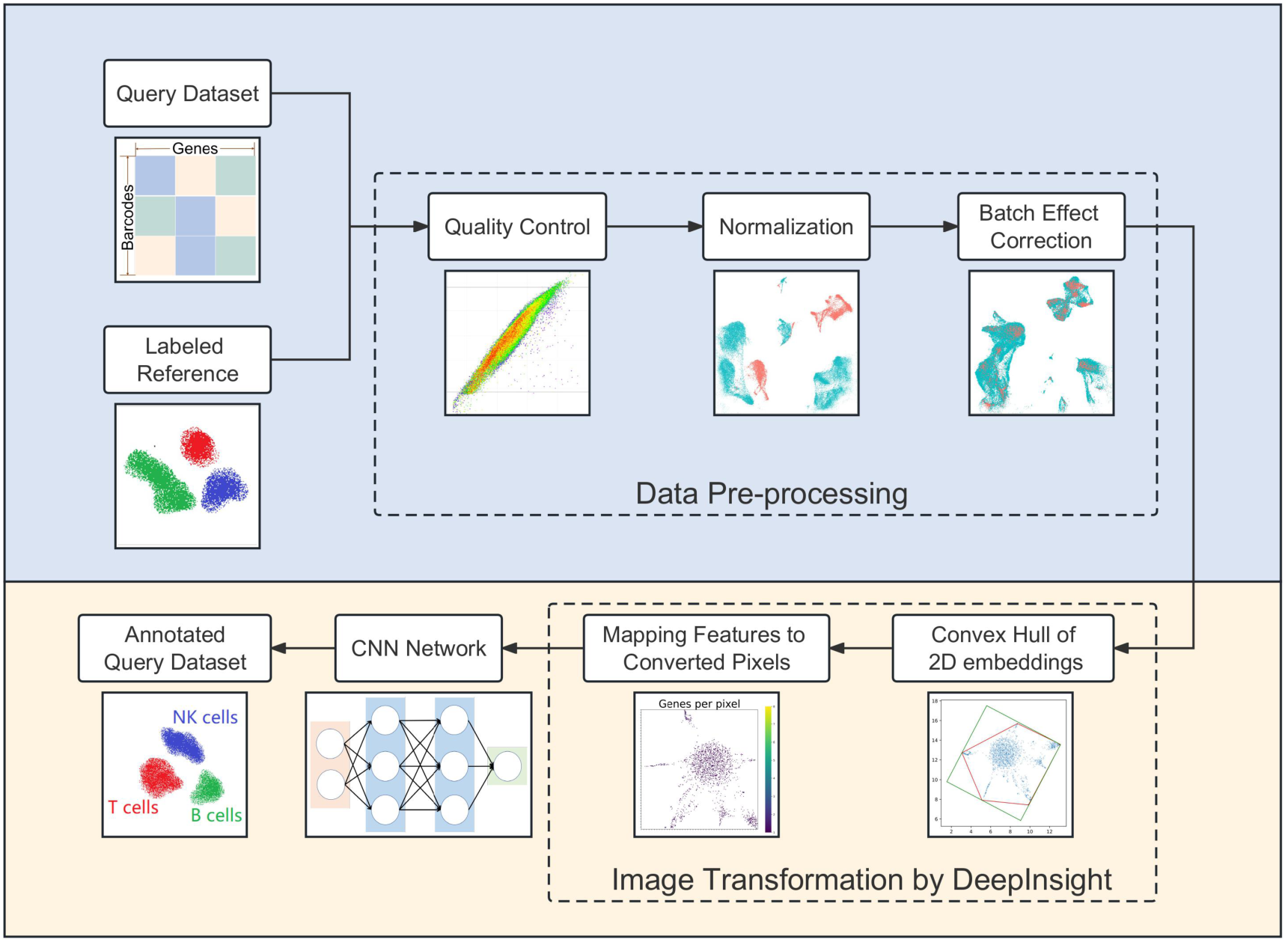
The scDeepInsight pipeline: the key steps performed by scDeepInsight from inputting single-unique molecular identifier count matrix to outputting cell annotation prediction. A reference dataset with a query data is processed via quality control, normalization and correction of batch effects. Then processed tabular data are converted into 2D embeddings. After framing and feature mapping, single-cell expression data are transformed into corresponding images. After this step, the reference dataset is used in training the CNN model. In the training step, no query dataset is used. Once the CNN model is trained, it is used to cluster single-cell samples from query dataset into cell-types. For subsequent query datasets, no further training of the reference dataset is performed, and therefore, the previously trained model can be directly used for clustering and annotation.

## Materials and Methods

### Overview of scDeepInsight

The workflow of scDeepInsight integrates the whole process of single-cell annotation, including data preprocessing, image conversion, neural network training and cell-type prediction (Figure 1). It takes individual unique molecular identifier (UMI) count matrices generated after sequencing as input. Each row of the matrix represents one cell with a unique barcode, and each column represents a gene. As scDeepInsight is a single-cell labeling model based on supervised learning a reference dataset is also required. After preparing reference and test datasets, data preprocessing is performed. This step includes quality control, normalization, and correction of batch effect between the query dataset and reference dataset. Afterwards, DeepInsight is utilized to convert the processed non-image data into images. First, the processed data are transformed into two-dimensional embeddings by some visualization method, like t-SNE. By mapping genes to pixels, the expression of different genes in a sample is transformed into a unique image. The lighter the pixel shade is in the image, the higher is the expression level of the gene in this cell. Next, the processed images from the reference dataset are used as the input for the CNN. After training, images transformed from the query dataset can be fed into the trained model to complete the annotation of cell-types. The output of the model is a vector of probabilities that a cell belongs to a specific cell-type. The label corresponding to the highest probability is taken as a cell-type prediction for that single-cell sample.

### Sample quality control

Quality control directly impacts the reliability of the downstream analysis. By controlling the number of specific genes detected in a single-cell (nFeature_RNA), the total number of UMI detected (nCount_RNA) and the proportion of mitochondrial genes (percent.mt) in each cell, cells with little effective information can be filtered out. Low-quality cells or empty droplets typically have very few genes detected. However, if there is an overlap where two or more cells are captured simultaneously, UMI detected in such cells will also become abnormally large. Both of these two cases should be removed. In this paper, we generally limit nFeature_RNA to between 300 and 4000 for single-cell samples. In addition, dying cells often exhibit extensive mitochondrial contamination. In this step, we set the threshold of percent.mt to 15 and cells exceeding this value are filtered out to avoid excessive influence of mitochondrial genes.

### Gene expression normalization

Commonly used normalization methods such as Scanpy::zheng17 [16] include selection of highly expressed genes, normalization, and scaling. In scDeepInsight, we use SCTransform [17] to normalize the expression data. This method performs regularized negative binomial regression on total UMI counts per cell to eliminate the variance due to sequencing depth. UMI reads are commonly positively correlated with the sequencing depth of the cells. Traditional logarithmic normalization process cannot remove this correlation among different cell samples. When extracting variable genes, genes selected by SCTransform also demonstrate more biologically meaningful variation than those chosen by the traditional normalization methods [18]. In addition, by performing regression on the percentage of mitochondrial genes or cell cycle of samples, SCTransform can also remove the influence of these factors.

### Batch Effect Correction

Batch effects have a quantitative impact on single-cell gene expression values. As a consequence, cells that should have been clustered together may end up divided into different clusters due to batch effects. We implemented several methods to eliminate the batch effect, including Canonical Correlation Analysis (CCA), ComBat and Harmony. Out of these methods, we found that CCA performed the best in most experiments. CCA is a multivariate statistical method to study the correlation between two groups of variables, and it can reveal internal relationships between variables. By calling Seurat::IntegrateData [19], the CCA method can be used to find anchors between datasets and integrate multi-sample datasets accordingly. This step can avoid the impact of biological heterogeneity caused by different experiment batches or sequencing technology on the accuracy of subsequent analysis.

### Data Scaling

The processed data needs to be scaled to the range [0, 1] before it is passed on to the image converter. For scRNA-seq data, two scaling methods are commonly used. The first is according to the maximum and minimum values of the expression data of each gene. The second is to take the dataset as a whole and scale it according to the global maximum and minimum values. In actual experiments, it appears that the values of some highly expressed genes are much higher than that of other genes. For this reason, the data processing function normalize_total of Scanpy also provides the option to ignore some particularly highly expressed genes when calculating regularization. In this context, differences in gene expression across cells are attenuated if scaled using global maximum and minimum. Therefore, each gene was scaled separately according to corresponding maximum and minimum values of expression.

### Generation of images from tabular data

We utilized DeepInsight [13] to map the high-dimensional feature space of scRNA-seq data into a 2D image space, where each pixel corresponds to a gene, and its intensity reflects the expression level of that gene in the cell. We used pyDeepInsight [https://github.com/alok-ai-lab/pyDeepInsight] to perform the image conversion with t-SNE to generate the 2D embeddings. To avoid overlapping of genes in the same pixel, we varied the perplexity value of t-SNE to identify the best-performing embedding using the training subset of the PBMC dataset (see Supplementary Figure 1 for different perplexity values) [20–22]. The 2D embeddings were subsequently transformed into images with the Cartesian plane for positioning features. This 2D positioning of features is determined by their similarity, and DeepInsight uses the convex hull algorithm to find the smallest rectangle containing all feature points and rotate it to the horizontal or vertical direction applicable to CNN architecture. Finally, Cartesian coordinates are converted to pixels according to the specified output image size, and the feature/gene values are mapped to these locations in the pixel frame. After this transformation, we used EfficientNet-b3 [23], a CNN pre-trained on large image datasets that can extract critical features that are integrated at the end for discriminating patterns, to analyze and classify scRNA-seq data.

### CNN model training and validation

To take advantage of the feature extraction ability and shorter training time achievable by using transfer learning, EfficientNet-b3 CNN model, which was pre-trained on large image datasets, was used for all analysis in this paper. Additionally, label smoothing and early stopping optimization techniques were used to reduce overfitting. All datasets used in the experiments were first subdivided into training and validation parts at 85:15 ratio. One of the accuracy plots during the training process is shown in Supplementary Figure 2.

## Results

### Datasets and preprocessing

The database of peripheral blood mononuclear cells (PBMC) offers readily accessible heterogeneous cell samples that contain several similar but distinct cell-types. PBMC datasets are usually dominated by several cell-types and are very unbalanced datasets. Furthermore, there are many similar cell subtypes, such as CD4+ Central Memory T (CD4 TCM) and CD4+ Central Effector T (CD4 TEM). Correctly annotating PBMC cells has always been a challenging task for researchers [7, 11]. In this paper, we used a PBMC dataset that had been labeled by experts in prior experiments [24] as a reference dataset for scDeepInsight training. The reference dataset contained over 160,000 single-cell samples and 31 different cell-types.

Independent test (or query) datasets used in this paper for benchmarking scDeepInsight are described in Table 1. The number of cell samples and cell-types contained in test datasets were relatively large. In addition to the five query datasets from the 10X Multiome sequencing protocol, we also used the data from the Wilk [25] study in our benchmarking analysis. The Wilk dataset was generated using seq-Well sequencing technology and contains 15765 cells. The dataset was preprocessed using the same pipeline as the 10X Multiome datasets, including quality control, gene filtering, and normalization. However, the nFeature_RNA and nCount_RNA of the Wilk dataset were much smaller than those of the reference dataset, with an average of 1852 genes detected per cell and an average of 6497 UMIs per cell (Supplementary Figure 3). Notably, the expression levels varied greatly between the reference and test sets. Despite the differences in sequencing protocols and data characteristics scDeepInsight outperformed other methods on this dataset as well, as shown in Figure 3. This suggests that scDeepInsight is robust to differences in sequencing protocols and data characteristics.

**Table 1.** Summary of inter-datasets.

| Dataset | Genes | Cells | Protocol | Cell types |
| --- | --- | --- | --- | --- |
| Hao (reference) | 20729 | 161764 | 10x Multiome 3' v3 | 31 |
| Yazar (test) | 36571 | 255734 | 10x Multiome 3' v3 | 25 |
| Schulte-Schrepping (test) | 33228 | 45787 | 10x Multiome 3' v2,v3 | 20 |
| Arunachalam (test) | 22878 | 49139 | 10x Multiome 3' v3 | 26 |
| Lee (test) | 22021 | 43512 | 10x Multiome 3' v3 | 24 |
| 10x-Multiome-Pbmc10k (test) | 29095 | 9631 | 10x Multiome | 19 |
| Wilk (test) | 24955 | 15765 | Seq-Well | 27 |

The test dataset needs to be preprocessed before using DeepInsight for image conversion. After quality control, cell samples with poor sequencing quality were filtered out (Figure 2a; see Materials and Methods section). Then we used SCTransform to complete the screening for highly expressed genes and do the normalization. Dealing with batch effects between reference and test datasets is an important step in preprocessing. Samples sequenced by 10x Multiome 3’ v3 in the test dataset used the same sequencing technology as the reference dataset, so the sequencing depth (total UMI counts per cell) was approximately the same. However, in the part of the test set sequenced by 10x Multiome v2 and Seq-Well, fewer genes and UMI counts were detected, and the sequencing depth was shallower than that of the reference dataset. Such differences due to sequencing platforms should be addressed when dealing with batch effects. Experiments on the test dataset Schulte-Schrepping [26] showed that CCA could successfully eliminate the batch effect (Figure 2b). Prior to batch effect correction distribution patterns displayed high degree of variation between the samples from reference and test datasets, even though these cell samples were of the same cell-type (such as CD4 TCM and CD4 TEM cells). After eliminating the batch effect, the distribution difference between expression data in UMAP results was mainly determined by cell-types, rather than the batch effect arising from the sequencing platform differences between the test and reference datasets (Figure 2c).

**Figure 2.**
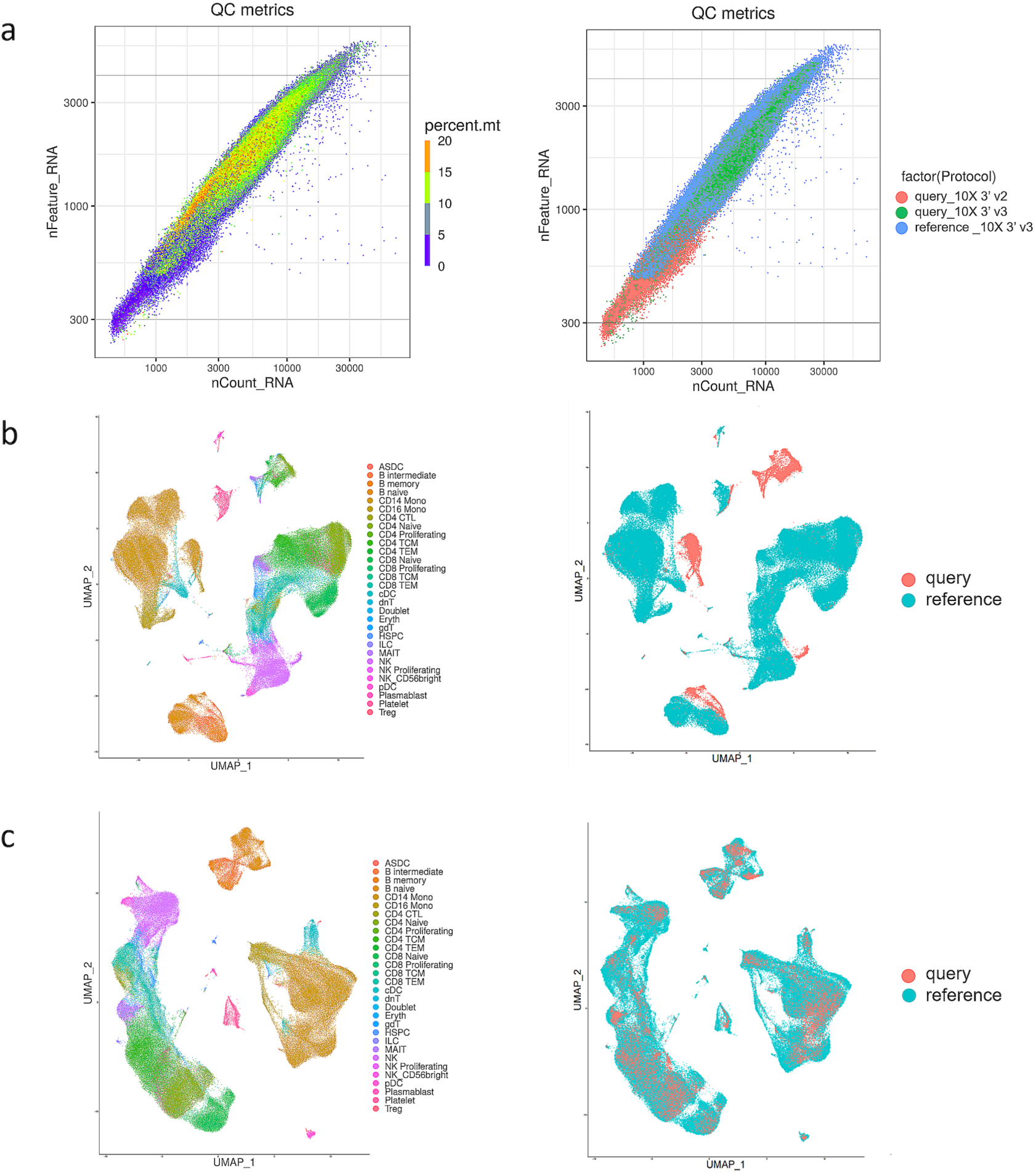
Preprocessing results of the test dataset Schulte-Schrepping. (a) The left plotis the quality control plot of the test dataset. Cells whose nFeature_RNA was less than 300 or more than 4000 were filtered out. Cells with percent.mt larger than 15 were also excluded. The right plot is labeled by the sequencing technology of data: samples in the reference dataset sequenced by 10x Multiome 3’ v3, samples in the query dataset sequenced by 10x Multiome 3’ v2 and 10x Multiome 3’ v3. (b) The Uniform Manifold Approximation and Projection (UMAP) representation of the reference before batch effect correction labeled by cell-types and data sources (c) The UMAP representation of datasets after batch effect correction.

### Parameter settings and intra-dataset analyses

We conducted several intra-dataset analyses with different perplexity parameters and distance functions for t-SNE-based image conversion of the PBMC reference dataset. The mappings between converted pixels and eigengenes generated by scDeepInsight are shown in Supplementary Figure 1a for various perplexity parameters. We also evaluated the classification accuracy of different perplexities using intra-dataset analysis on the PBMC reference dataset, and the results are presented in Supplementary Figure 1b. Additionally, we employed a Bayesian optimization technique that takes into account the specifics of the training dataset, allowing us to search for the most appropriate hyperparameters. Supplementary File 1 provides reasonable ranges for the initial and final values selected by the optimization algorithm for the training datasets.

To further demonstrate the robustness of scDeepInsight, we performed intra-dataset analysis with scDeepInsight on datasets from other tissues. Specifically, we analyzed the kidney dataset (https://doi.org/10.48698/3z31-8924) from the Kidney Precision Medicine Project (KPMP) and pancreas dataset Tabula [22] generated using Smart-seq2 protocol. The kidney dataset created by KPMP contains 107344 cells from the human kidney, and Tabula dataset contains 3384 cells from the mouse pancreas. Both datasets were preprocessed using the same pipeline as the one used for the 10X Multiome datasets, and scDeepInsight was able to successfully eliminate batch effects on these datasets as well, as shown in Supplementary Figure 4. For the kidney dataset, the annotation accuracy on the test set by scDeepInsight was 93.8%, whereas the second best performing method, singleR, scored 86.7%. For the pancreas dataset Tabula generated using Smart-seq2 protocol, the accuracy achieved by scDeepInsight was 95.8%, and second-best method scBert had 89.2% annotation accuracy.

### Performance of scDeepInsight with inter-datasets and comparison with other methods

Inter-dataset accuracy was measured on six independent PBMC test datasets with high-quality pre-annotated labels. We used accuracy and adjusted Rand index (ARI) to measure the quality of cell-type predictions. Accuracy represents the percentage of predicted cell-types that perfectly fit with the cell-types labeled in previous reliable experiments. ARI is a measure of similarity between real cell labels and predicted cell-type clusters. The higher the value of ARI, the more similar the predicted results are to the original labels of cell samples. Moreover, to provide a more comprehensive evaluation of our method’s performance, we have included additional metrics such as F1-score and AUROC in addition to accuracy. As the PBMC datasets used in our study are imbalanced, we have also included AUPRC metric in our benchmarking process. All of these results are presented in Supplementary Figure 5. Our findings demonstrate that scDeepInsight outperforms other methods not only in accuracy but also in F1-score, AUROC and AUPRC, indicating the robustness of our approach in identifying cell-types from scRNA-seq data.

To benchmark the accuracy of our method, we have compared its performance to different kinds of mainstream cell annotation methods. The first type was annotation based on marker genes, including SCINA, SC3 and Seurat::FindClusters. As a graph-based clustering algorithm provided by Seurat, FindClusters is able to identify clusters through shared nearest neighbor (SNN) modularity optimization. Unlike SC3 and SNN, SCINA does not use unsupervised clustering. Instead, SCINA directly performs enrichment analysis on the specific marker gene list to assign type annotation. Furthermore, we chose ScType as a marker gene database. Integrating two marker gene databases, CellMarker and PanglaoDB, it provides a comprehensive database of specific markers covering many cell-types. In this paper, we refer to marker genes of the immune system in the ScType database to complete the annotation of clustering results. In order to ensure the fairness of the comparison, the marker genes were not manually selected when implementing these methods. For the reference-based annotation method, we chose SingleR, which achieves cell-type annotation by calculating and comparing the Spearman correlation coefficient between single-cell samples in query and reference datasets. To guarantee the validity of benchmarking, the same reference dataset used in scDeepInsight was utilised for learning. We also tested CellTypist and scBERT to make the comparison more comprehensive. Being an automatic labeling method, CellTypist offers built-in, pre-trained models for different tissues by integrating multiple single-cell data sets. The model we chose was Healthy_COVID19_PBMC, which best fitted the test datasets. Similarly, scBERT also provides a pre-trained model trained on a large-scale reference dataset PanglaoDB [6] and can perform cell-type annotation after fine-tuning the model with the target dataset. When implementing other annotation methods in the benchmarking process, we utilized the parameter values recommended by the original authors and performed the standard pipelines used by them.

On six test datasets, our method outperformed the other six methods (Figure 3 and Supplementary Figure 5). The average accuracy of scDeepInsight (87.5%) was more than 7% greater than the next best performing method, singleR. Specifically, on the dataset Yazar, the accuracy was the highest (96.1%). Also, the average ARI of scDeepInsight 0.851 on these six datasets was about 0.07 higher than that of other methods, which shows that the prediction results of scDeepInsight could better reflect real clustering patterns of cell labels. Other metrics also confirmed the superiority of scDeepInsight (Supplementary Figure 5). The average of F1-score, AUROC, AUPRC improved by 0.092, 0.109 and 0.105, respectively, relative to the next-best method. In addition, these query datasets included both 10X Multiome 3’v2, v3 and Seq-Well sequencing technologies. Despite the differences in sequencing protocols and data characteristics scDeepInsight outperformed other methods on this dataset as well, as shown in Figure 3 and Supplementary Figure 5. This suggests that scDeepInsight is robust to differences in sequencing protocols and data characteristics.

**Figure 3.**
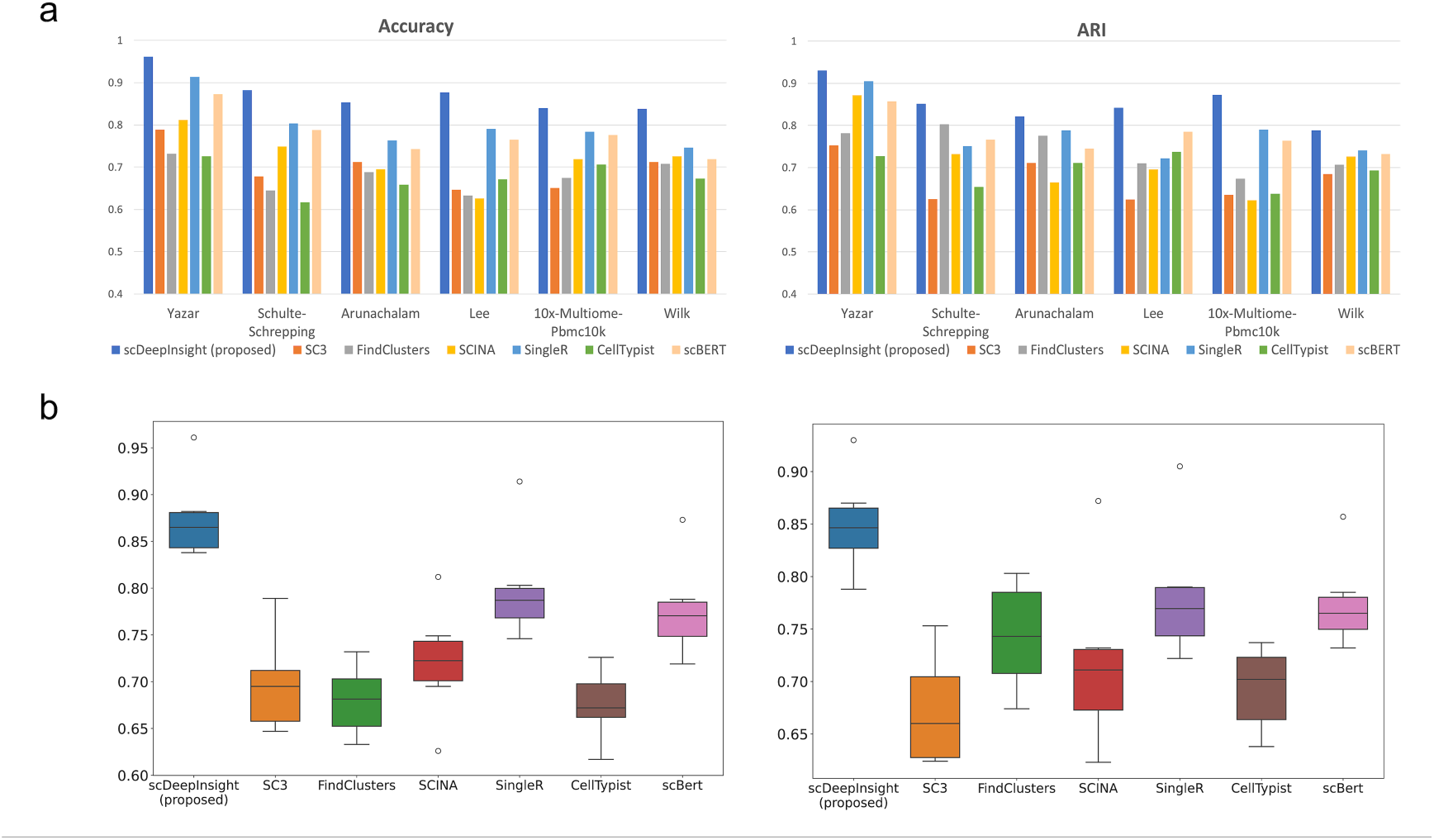
The performance of scDeepInsight. (a) Accuracy and ARI of scDeepInsight compared to other methods: SC3, FindClusters, SCINA, SingleR, CellTypist and scBERT, across the six datasets: Yaza, Schulte-Schrepping, Arunachalam, Lee, 10x-Multiome-Pbmc10k and Wilk. (b) The accuracy and ARI box plots of scDeepInsight and the other six methods used in benchmarking are depicted.

Furthermore, scDeepInsight had a high recognition accuracy for main cell-types in the test dataset (Figure 4). As shown in Figure 4a, 99.4% of CD14 Monocytes were correctly labeled as CD14 Mono by scDeepInsight. CD14 Monocytes accounted for more than 35% of the test dataset (Figure 4b). The heatmap of the confusion matrix shows that high recognition accuracy was achieved for most cell-types (Figure 4c; corresponding detailed confusion matrix is shown in Supplementary Figure 6). The cell-type prediction results by scDeepInsight were largely consistent with the original labels and further subdivided into some cell-types (Figure 4d).

**Figure 4.**
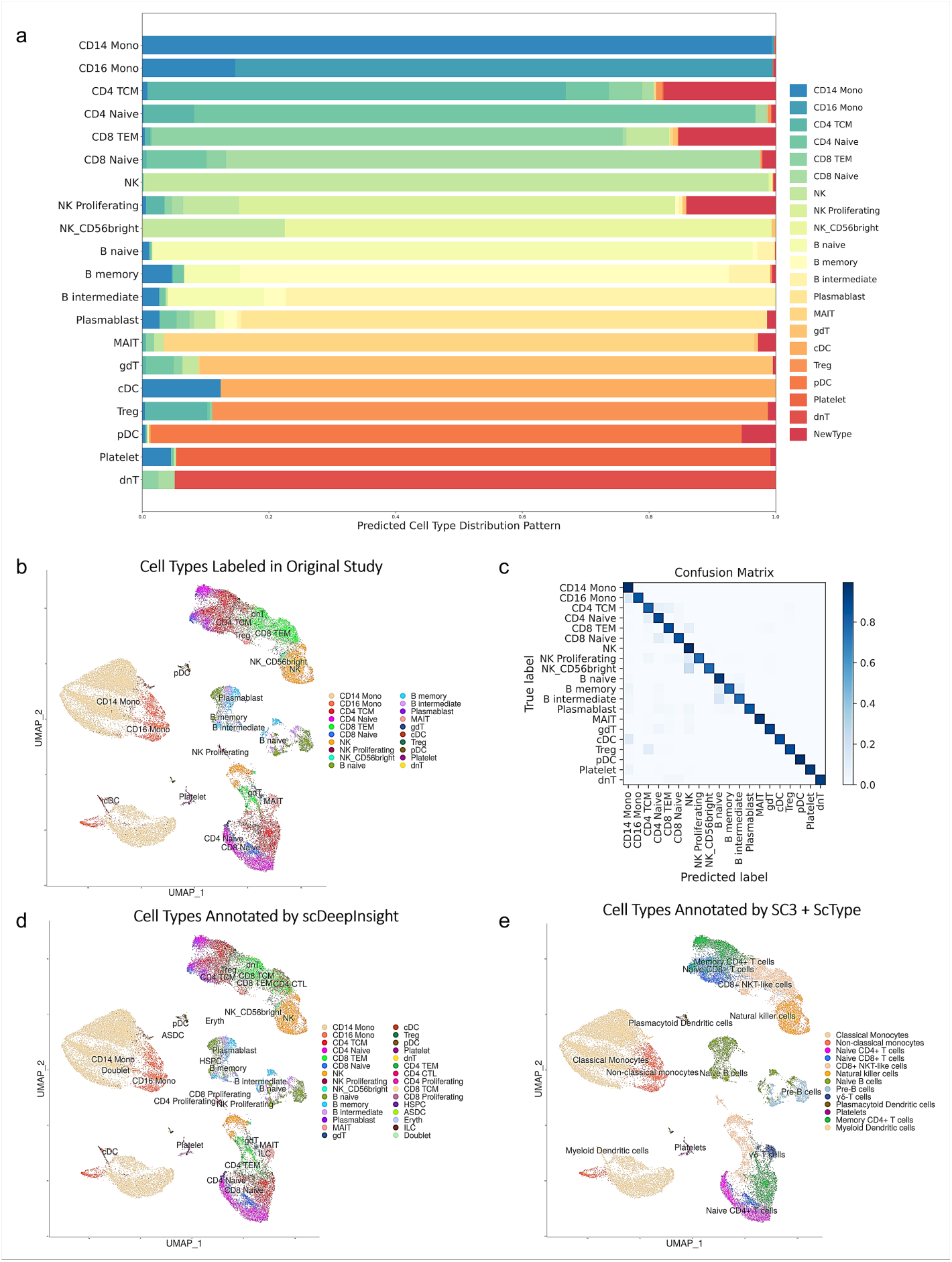
Cell-type labels of the reference dataset and prediction results on dataset Schulte-Schrepping. (a) The stacked percentage column chart of the prediction results on the Schulte-Schrepping dataset. (b) UMAP representation colored by cell-types in the original study (c) Heatmap of the confusion matrix. (d) UMAP representation colored by cell-types predicted by scDeepInsight. (e) UMAP representation colored by annotation results using SC3 clustering with ScType.

The other unsupervised clustering methods rarely found the correct number of clusters. Some of the cell subtypes, including CD4 Proliferating and CD8 Proliferating, could not be recognized using other annotation methods. Cell-type annotation results of the Schulte-Schrepping dataset using SC3 with ScType are shown in Figure 4e. Samples were first clustered by SC3 and then annotated by referring to the marker gene database ScType. Compared with prediction results by scDeepInsight, this annotation method recognized fewer clusters and cell subtypes.

In conclusion, compared with other mainstream sequencing methods, our method was not only more accurate but also more capable of detecting similar cell subtypes.

### The ability to discover new cell-types

The ability to discover new cell types is a critical feature of scDeepInsight that distinguishes it from many existing methods for cell-type annotation. Traditional annotation methods rely on reference datasets or marker gene databases, which may not cover all cell-types present in a given test dataset. Unsupervised methods can reveal new cell-types by detecting clusters that cannot be annotated to known cell-types, but many supervised methods simply label all samples as known cell-types contained in the reference dataset. This approach can ignore rare or unknown cell-types, which can significantly impact the accuracy of cell-type labeling.

scDeepInsight addresses this limitation by returning the probability that each single cell should be labeled as a specific cell-type. By setting a probability threshold we can filter out the cases where the predicted probability for all cell-types is very low, indicating that the sample is not similar to any known cell-types in the reference dataset. In such cases, these cells can be predicted as an unknown type.

To demonstrate scDeepInsight’s ability to identify rare or unknown cell types, we evaluated its performance on two datasets containing cells from COVID-19 infected patients: the Lee [27] and Arunachalam [28] datasets. These datasets contain some neutrophils, activated natural killer cells, activated CD4-positive T cells, and activated CD8-positive T cells, which do not exist in the reference PBMC dataset derived entirely from healthy donors. We found that the predicted probabilities returned by the fully connected layer for cells from healthy donors and infected patients are not identical. Different distribution plots of predicted probabilities are depicted in Supplementary Figure 7. When filtering out 1% of cells with the smallest predicted probability, 72 out of a total of 89 neutrophils contained in the Lee dataset can be correctly detected and annotated as unknown cell-types. For the Arunachalam dataset, 1536 out of a total of 1880 activated immune cells not contained in the reference dataset can be successfully identified under the same threshold condition.

Identifying rare or unknown cell types has important implications for our understanding of disease processes and the development of new treatments. For example, the identification of new immune cell subtypes could lead to the discovery of new therapeutic targets or improve our understanding of how the immune system responds to viral infections. In summary, scDeepInsight’s ability to identify rare or unknown cell types is a valuable feature that has the potential to significantly advance our understanding of complex biological systems.

### Identification of marker genes

By analyzing the fully connected layer in a trained CNN, DeepFeature [15] can construct Class Activation Mapping (CAM) to extract features of different classes. In this study of the single-cell classification, we also introduced CAM to help analyze the features differentially expressed among multiple cell-types. After applying DeepFeature to single-cell datasets, it was found that the genes extracted in different cell-types contained marker genes corresponding to these types. Generated CAM graphs are shown in Supplementary Figure 8. Through the analysis of the extracted marker genes, the accuracy of the cell-type annotation model established by scDeepInsight at the biological level could also be proven. We selected three marker genes for Monocytes extracted by scDeepInsight from the reference dataset and colored the UMAP representation of the reference according to the expression values (Figure 5a). By referring to the cell-type distribution of the reference dataset shown in Figure 5b, the differential expression of these three genes in different regions also confirmed their reliability as marker genes.

**Figure 5.**
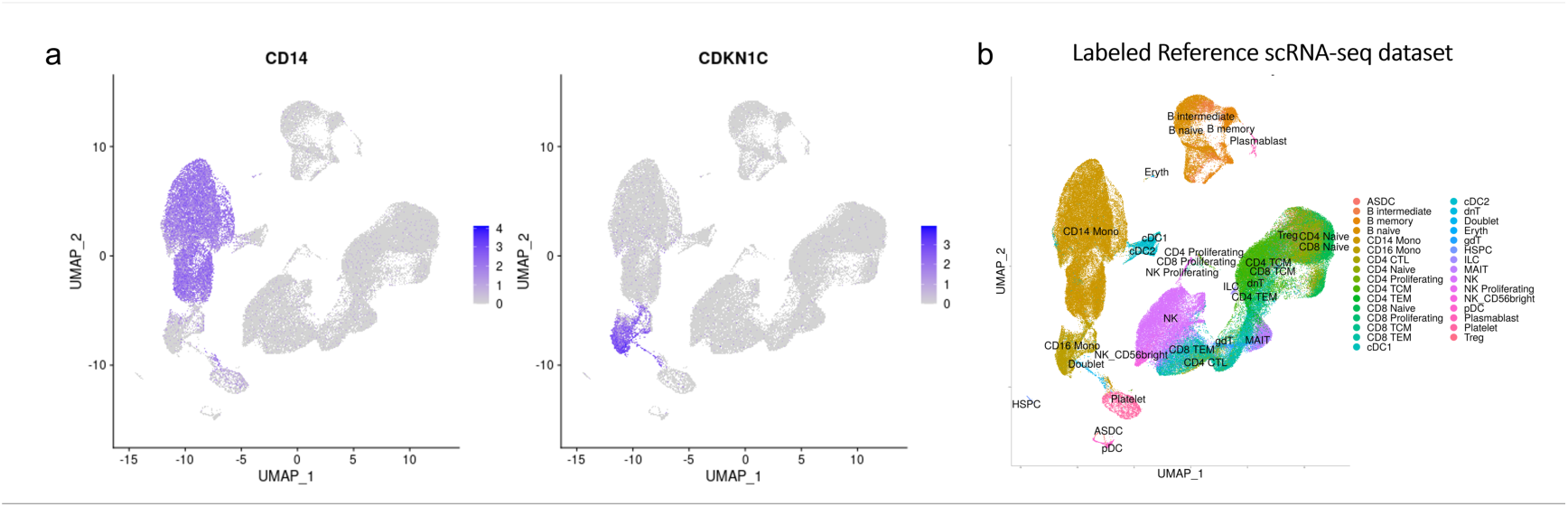
(a) CD14 [29] and CDKN1C [30] were proven to be marker genes for monocytes, CD14 Monocytes and CD16 Monocytes correspondingly in the previous study. (b) The UMAP 2D embedding after performing normalization and principal component analysis (PCA) dimensionality reduction on the reference dataset. Cells are grouped and colored by known labels from previous studies.

### Summary of overall performance

Compared with unsupervised clustering algorithms, supervised labeling methods based on reference datasets can often achieve more accurate identification. However, supervised annotation methods require a correctly labeled reference dataset as the prerequisite, which is a limitation. In addition, training on reference datasets also brings additional time costs. Nonetheless, as an original supervised cell-type annotation method, scDeepInsight converts preprocessed single-cell data into images and utilizes CNN’s strengths in feature extraction and classification, enabling accurate identification of cells in a few epochs. We have shared our pretrained models so that other researchers can easily use scDeepInsight to annotate their own datasets. On large-scale PBMC test datasets, both annotation accuracy, ARI and AUROC have improved relative to other mainstream methods. Furthermore, scDeepInsight is capable of detecting new cell types not contained in the reference dataset. With the assistance of DeepFeature, our method can also extract marker genes for multiple cell types.

## Discussion

DeepInsight, on which scDeepInsight is based, has several advantages, including its flexibility, ability to reveal patterns and insights that are not easily observable in the original feature space, and interpretability of results through visualized features. Its approach of positioning features in the Cartesian plane instead of samples (i.e. cells) allows it to avoid dimensionality reduction of the features [20]. Also, it can utilize various visualization techniques, such as PCA, UMAP, or kernel PCA, as alternatives to t-SNE, which we have employed in this work due to its high performance. Kobak and Berens propose an improved protocol for visualizing single-cell transcriptomics data using t-SNE that includes PCA initialization, high learning rate, and multi-scale similarity kernels [21]. They recommend exaggeration and downsampling-based initialization for large datasets, and demonstrate that this protocol yields superior results compared to naive application of t-SNE. They also discuss the advantages and disadvantages of t-SNE compared to UMAP, and describe how to position new cells on an existing t-SNE reference atlas and visualize multiple related data sets consistently. In relevance to the interpretability of results, we used CAM in DeepFeature [15]. Another option may be to use a “relevance aggregation” method that improves the interpretability of neural networks for tabular data [22]. The algorithm generates scores for each input feature by combining the relevance computed from several samples, making it easier to identify important features.

Although DeepInsight has several advantages, its limitation lies in the potential for genes to overlap in the same pixel due to the limited pixel frame size of the image. To avoid overlapping of genes in the same pixel, we set an appropriate resolution for the converted image. Wattenberg and colleagues [20] discuss the usefulness of t-SNE in visualizing high-dimensional data. They explained the importance of the perplexity parameter in creating accurate t-SNE plots and cautioned that the cluster sizes shown in these plots may not accurately represent the original cluster sizes in high-dimensional space. Its effectiveness relies on selecting parameters carefully and interpreting the plots thoughtfully. Therefore, when selecting t-SNE, we optimized and manually set the parameters for perplexity and distance function, with larger perplexity values recommended for datasets with larger numbers of cells. This approach not only helps to reduce obscurity caused by overlapping but also facilitates the identification of marker genes for specific cell types. To mitigate the stochasticity of t-SNE, another approach is to generate multiple embeddings using different random seeds and select the one with the lowest Kullback-Leibler divergence from the original data distribution. Alternatively, one can fix the random seed to obtain consistent embeddings for a particular dataset.

### Ablation study for multiple conditions

To further investigate the impact of individual steps in the pipeline on the accuracy of cell type annotations, we conducted ablation experiments under multiple conditions. In the data preprocessing stage, we compared various single-cell processing methods and tested whether data normalization and batch effect correction processes improved the accuracy of cell type annotation. As shown in Supplementary Figure 9a, batch effect correction increased accuracy by an average of 3.18% across six test datasets, while data normalization increased accuracy by an average of 0.82%. In the image conversion step, we replaced DeepInsight with numpy.resize to convert single-cell expression data into three-channel images and evaluated the resulting accuracy. The accuracy plot and the converted image are presented in Supplementary Figure 9b and 9c, respectively. Without DeepInsight’s image conversion the average accuracy dropped sharply to 15.2% on the six test datasets. However, using the scDeepInsight annotation pipeline with the same network structure and parameters, the average annotation accuracy rate can reach 87.5% on those test datasets.

Next we measured the choice of CNN models in extracting image features through convolution and pooling operations, which are then processed in the fully connected layer for image classification. Different CNN models have been developed based on differences in network structure and connection mode. scDeepInsight provides support for multiple CNN models, such as ResNet [31] and DenseNet [32]. In our ablation study, we compared the performance of various CNN models and found that EfficientNet-b3 [23] exhibited high efficiency and accuracy. These results are presented in Supplementary Figure 9. Moreover, scDeepInsight shows strong performance on multiple CNN networks, suggesting that changing the type of image network may not significantly impact recognition accuracy.

### Speed of scDeepInsight

Although the running time is hardware-dependent, our proposed scDeepInsight method can complete the annotation process of 10,000 cell samples within 60 minutes with one Intel(R) Xeon(R) Gold 6242 CPU (16 cores, 2.80GHz). Table 2 presents the processing time of various methods. Of the total time cost, 55 minutes are devoted to performing batch effect correction, while 5 minutes are used for image conversion and loading the pretrained model. If the batch effect correction method Seurat::IntegrateData is not used, and only basic preprocessing methods are applied to the target annotation dataset, the entire annotation process will be less than 3 minutes. However, using batch effect correction can increase the accuracy of the model, but also significantly increase the total time cost. For this reason, we provide details of the two annotation pipelines (with and without batch effect correction) and their corresponding pre-trained models on our GitHub repository. This way, users can choose whether to perform batch correction on the target data, based on their actual application scenario. Furthermore, as a supervised annotation method, scDeepInsight is efficient during the training process. The accuracy of the EfficientNet-b3 model on the validation set can converge to more than 95% within 50 epochs. When using a batch size of 128, the time cost for one batch is about 10 minutes using two GPUs (Quadro RTX 8000 48GB) with one Intel(R) Xeon(R) Gold 6242 CPU (16 cores, 2.80GHz).

**Table 2.** Execution time cost by different methods to annotate a dataset of 10,000 cells.

| Machine learning methods | Time cost (seconds) |
| --- | --- |
| scDeepInsight | 3327 |
| scDeepInsight (no batch correction) | 68 |
| SC3 | 341 |
| FindClusters | 196 |
| SCINA | 103 |
| SingleR | 3524 |
| CellTypist | 178 |
| scBERT | 1567 |

### Obtaining reliable reference datasets

We have provided guidance on how to select good reference data for use with scDeepInsight. Specifically, we recommend the use of reference datasets from Azimuth (https://azimuth.hubmapconsortium.org/references/) released by Satija lab, which have been widely used in the single-cell analysis community and have been shown to be reliable.

We also acknowledge that the availability of relevant reference datasets is still limited and that efforts to expand coverage of cell types that can be identified using supervised annotation methods are needed. Such datasets need to be produced experimentally with techniques such as FACS, sometimes accompanied with *in silico* insights for finding, defining, and accurately annotating cell types. One highly promising project in this regard is The Human Cell Atlas [33], an international collaboration aimed at mapping all cell types in the human body.

### Conclusions and future direction

In this paper, we proposed scDeepInsight, which integrates the whole operation of data preprocessing, image conversion and constructing the CNN model for cell-type prediction. The prediction results correctly reflected the number of clusters and further enabled outlier detection for unknown types. First, we selected a PBMC dataset containing trusted cell-type labels as a reference. By performing standard tests on six different independent test sets, it was shown that scDeepInsight had significantly higher accuracy and could identify more cell subtypes than the other six mainstream cell labeling methods. In addition, we addressed classical problems in single-cell annotation, such as batch effect correction methods and the detection of rare/unknown cell-types. Finally, by applying DeepFeature to extract marker genes of cell-types, the accuracy of scDeepInsight in feature extraction was further proved in biological significance.

In the future, we will integrate more reference datasets obtained from different tissues to construct a more comprehensive cell-type classification model. Also, we will try to integrate protein and spatial chromatin accessibility information to further improve the accuracy of cell-type identification. Furthermore, in subsequent versions of our software, we plan to provide pre-trained models based on reference datasets from different types of tissues to address the issue of limited availability of reference datasets.

## Data Availability

All datasets used in this paper are publicly available. The reference PBMC dataset [24] (GSE164378) can be obtained from [https://atlas.fredhutch.org/nygc/multimodal-pbmc/]. Dataset 10x-Multiome-Pbmc10k is provided by scglue [11] and can be downloaded directly from [https://scglue.readthedocs.io/en/latest/data.html]. Query datasets Yazar [34] (GSE196830), Schulte-Schrepping [26] (EGAS00001004571), Lee [27] (GSE149689), Arunachalam [28] (GSE155673),Wilk [25] (GSE150728), kidney dataset from KPMP (GSE183279) and Tabula [22] (GSE132042) are all available from the CELLxGENE database: [https://cellxgene.cziscience.com/datasets].

## Code Availability

scDeepInsight is available as a Python package, the latest version can be downloaded from the Python Package Index (PyPI): [https://pypi.org/project/scdeepinsight/]. In addition, to facilitate and simplify the deployment process, we have uploaded a ready-to-use environment: [https://hub.docker.com/r/shangrujia/scdeepinsight]. We also provide pretrained annotation models to save users time. The entire code base, including the implementation of the proposed scDeepInsight pipeline and the pretrained models, is available at: [https://github.com/shangruJia/scDeepInsight]. Another GitHub repository [https://github.com/shangruJia/scDeepInsight-additional] stores codes for reproducing the training and testing results used in the paper for benchmarking. Gene IDs and barcodes of the cell samples used in this paper are recorded in Supplementary File 2, which can also be downloaded from this repository.

## Acknowledgements

The results shown in this paper are in part based upon PBMC dataset generated by New York Genome Center (NYGC). We appreciate Gene Expression Omnibus (GEO) and European Genome-phenome Archive (EGA) providing access to the query datasets.

## Author contributions

SJ implemented the whole pipeline, evaluated the performance, and wrote the first draft and contributed in the subsequent versions of the manuscript. AL advised for the model and the evaluation, contributed in the manuscript writeups. KAB checked the model, and helped in the manuscript writeup. AS perceived, supervised, and contributed in the manuscript writeups. TT perceived, supervised, and contributed in the manuscript writeups. All authors read and approved the manuscript.

