## Supplement File 1 for "scDeepInsight: a supervised cell-type identification method for scRNA-seq data with deep learning"

^5^ Co-last authors

**Supplementary File 1**

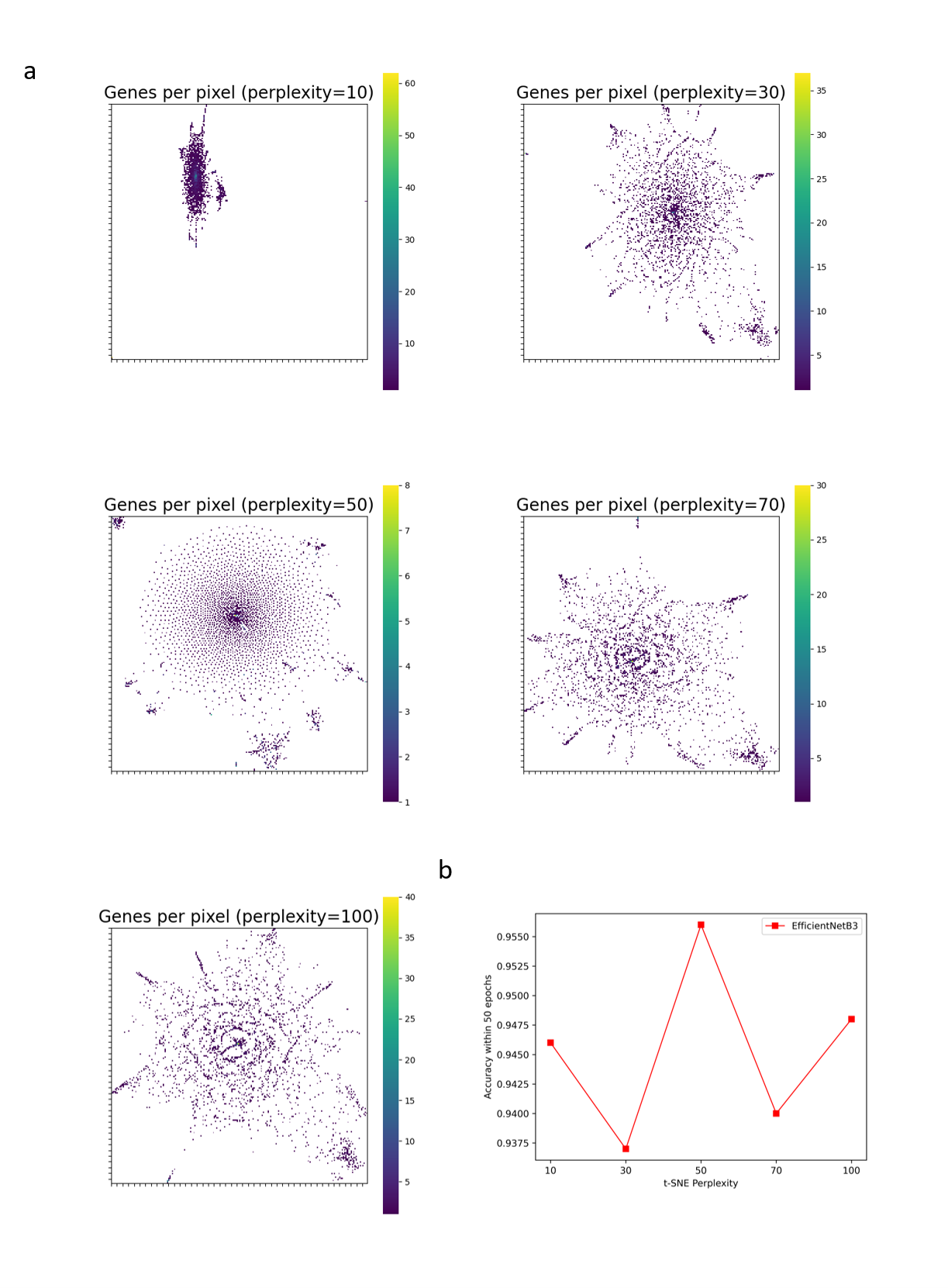

**Supplementary Figure 1. The effect of different t-SNE perplexity values tested in intra-dataset analysis optimization.** (a) Mappings between converted pixels and eigengenes generated by scDeepInsight under the perplexity of 10, 30, 50, 70, 100. (b) The cell-type annotation accuracy plot within 50 epochs plot in the intra-dataset tests. A perplexity value of 50 performed best and achieved an accuracy of 95.6% on the test set of the intra-dataset.

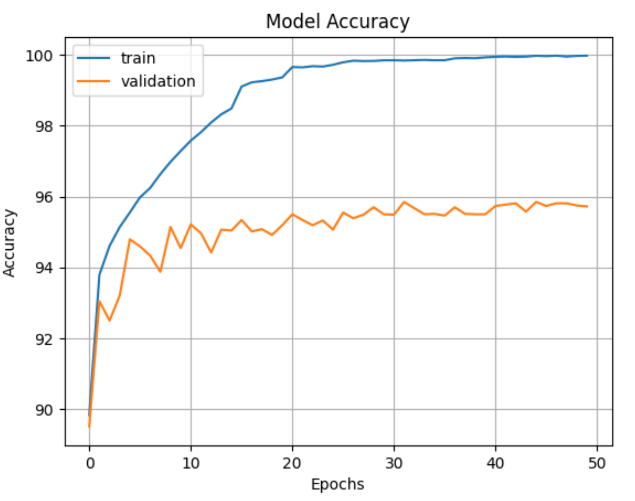

**Supplementary Figure 2. The trends of train accuracy and validation accuracy in the training process of the Lee dataset using EfficientNet-B3.** The size of input images was 224$\times$224 and the miniBatchSize was 128. The InitialLearningRate was 3$\times$10-4. To avoid overfitting, we used L2 regularization with a weight_decay parameter of 1$\times$10-6. In addition, we also introduced label smoothing to improve the generalization ability of prediction with a smoothing rate of 0.1. The validation accuracy reached 95.9% within 50 epochs.

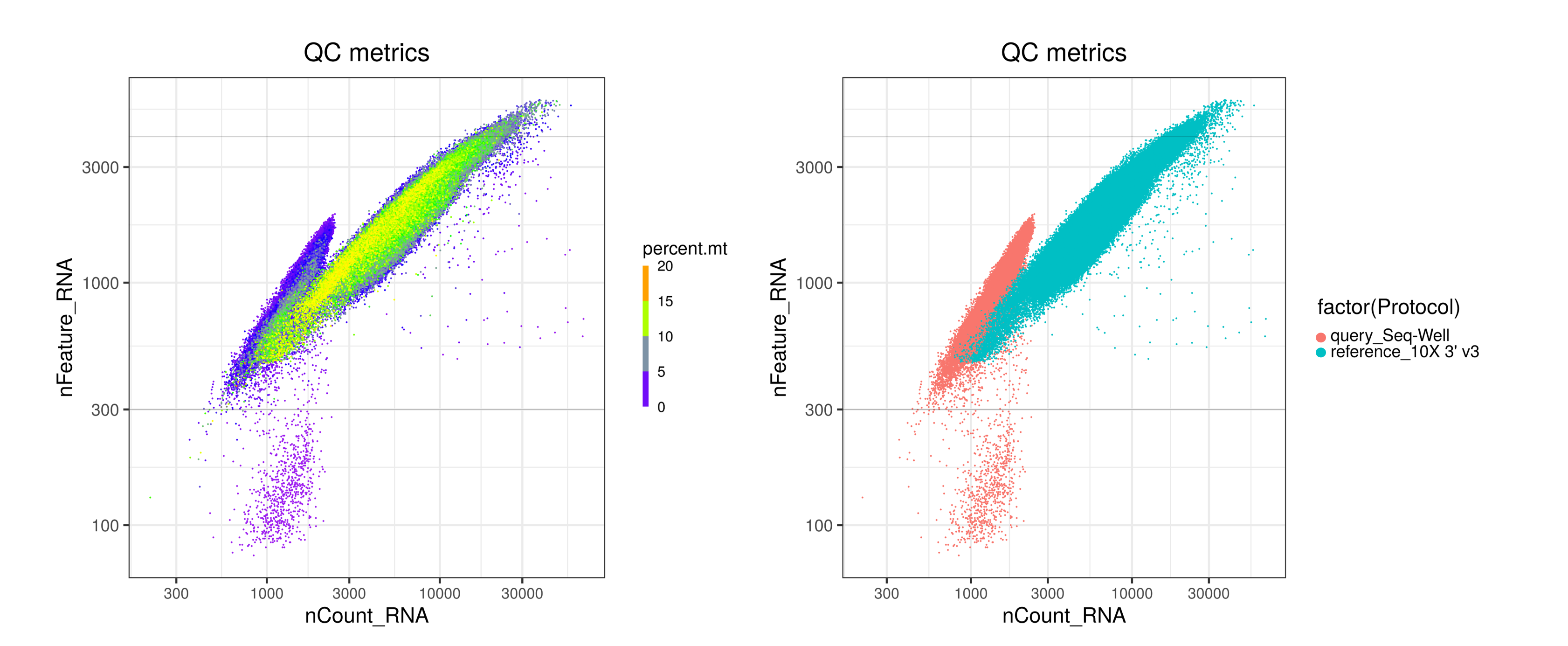

**Supplementary Figure 3. The quality control plot on the query dataset Wilk, labeled by percent.mt and sequencing protocols.** The left plot shows several metrics in quality control: the number of specific genes detected in a single cell (nFeature_RNA), the total number of UMIs detected (nCount_RNA) and the proportion of mitochondrial genes (percent.mt). The right plot is labeled by the sequencing protocol of datasets. For the test dataset Wilk, the sequencing method is Seq-Well, which is different from the reference dataset. Much fewer genes and a number of UMIs are detected using this sequencing technology, hindering the further integration of datasets. In such situations, batch effects between the test dataset and reference dataset bring the biological difference. The lack of effective gene expression information brought about by too shallow sequencing depth is difficult to be resolved in the batch correction step.

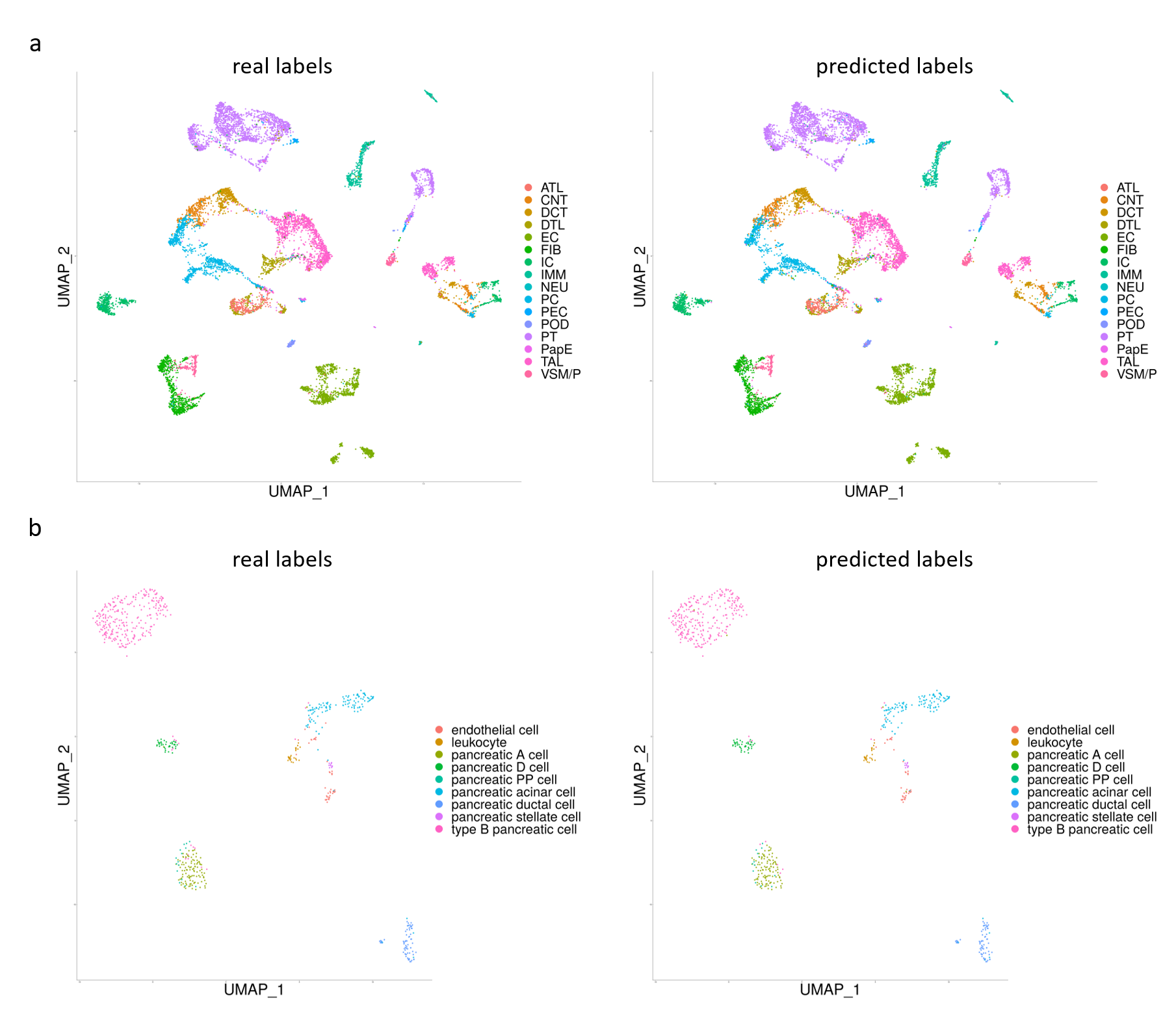

**Supplementary Figure 4. Real and predicted cell type distribution patterns of datasets from other tissues in the intra-dataset analysis.** (a) The left panel is the UMAP representation colored by cell-labels in the original study of the human kidney dataset from the Kidney Precision Medicine Project (KPMP). The right panel is colored by cell types predicted by scDeepInsight. (b) The left panel is the UMAP representation colored by cell-labels in the original study of the Tabula dataset from mouse pancreas. The right panel is colored by cell types predicted by scDeepInsight.

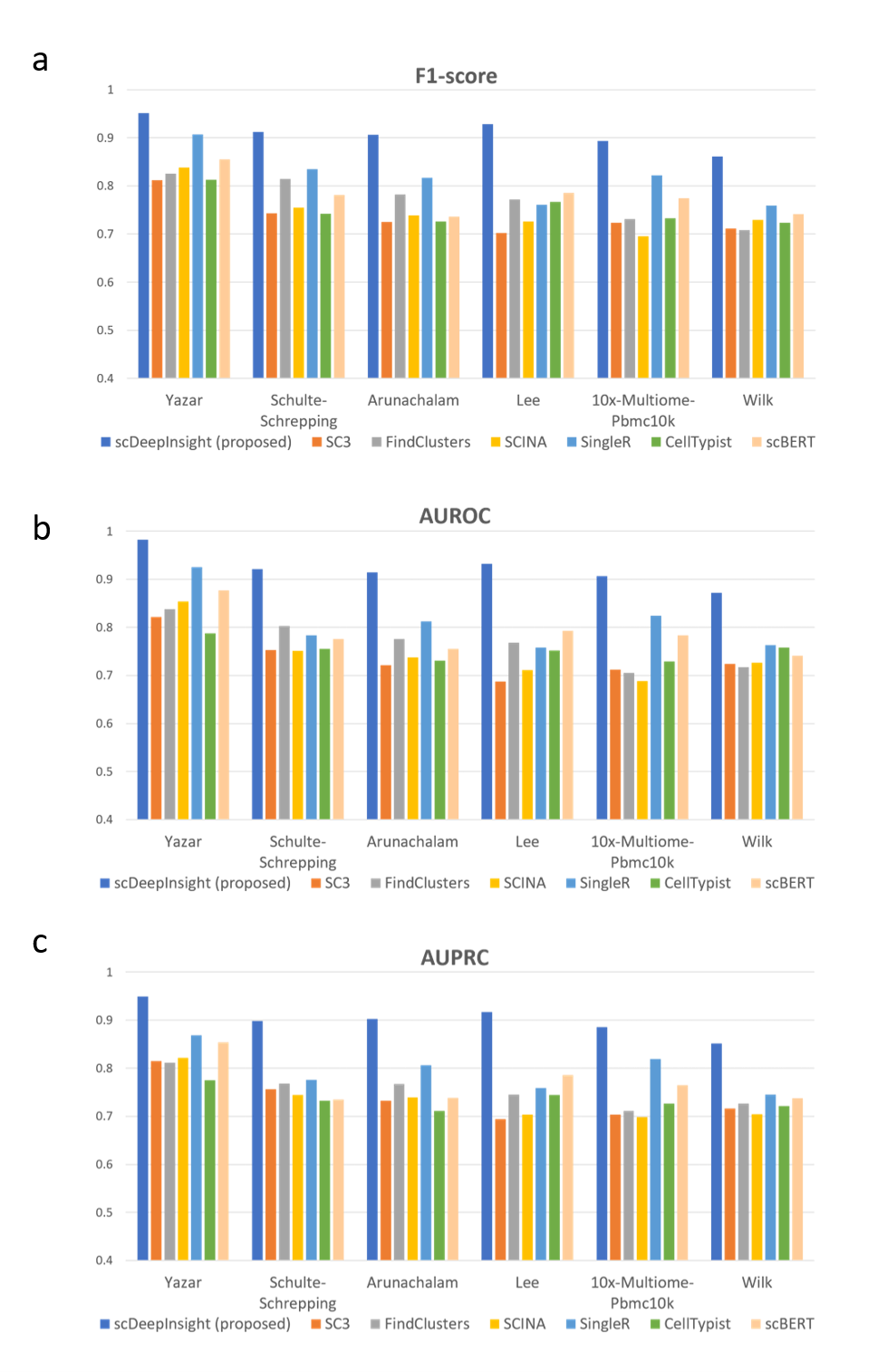

**Supplementary Figure 5. Performance evaluation of scDeepInsight on six inter-datasets with additional metrics.** The average (a) F1-score, (b) AUROC, and (c) AUPRC improved by 0.092, 0.109 and 0.105, respectively, compared with the second-best method, singleR.

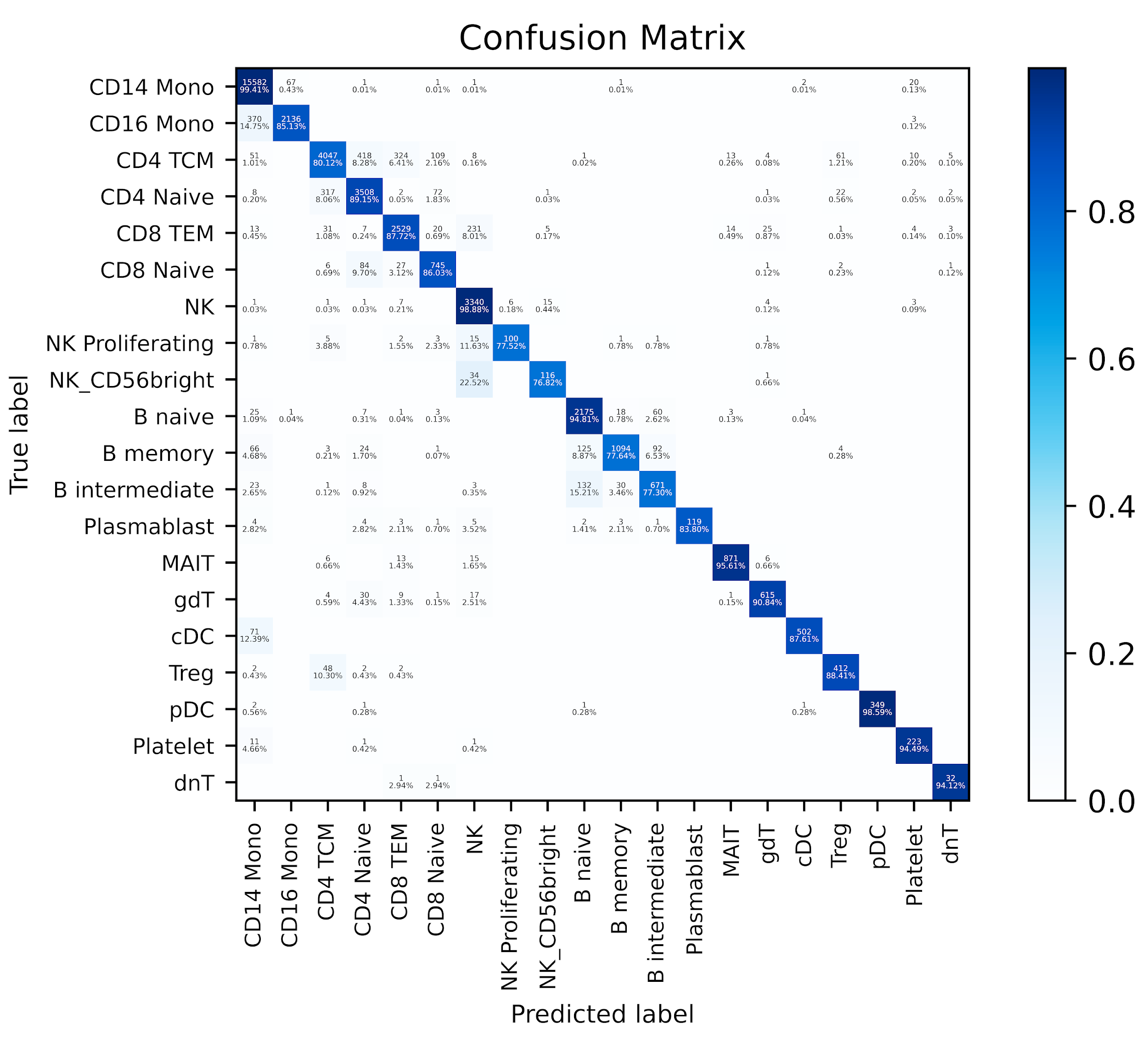

**Supplementary Figure 6. The heatmap for confusion matrix of the prediction result on the Schulte-Schrepping dataset.** For CD14 Monocytes, which account for the most dominant type in the test set, the classification accuracy can reach 99.41%. Natural killer (NK) cells, plasmacytoid dendritic cells (pDCs) and B naive cells are the other three cell-types with the highest accuracy. The overall accuracy on these 20 cell-types is 92.1%.

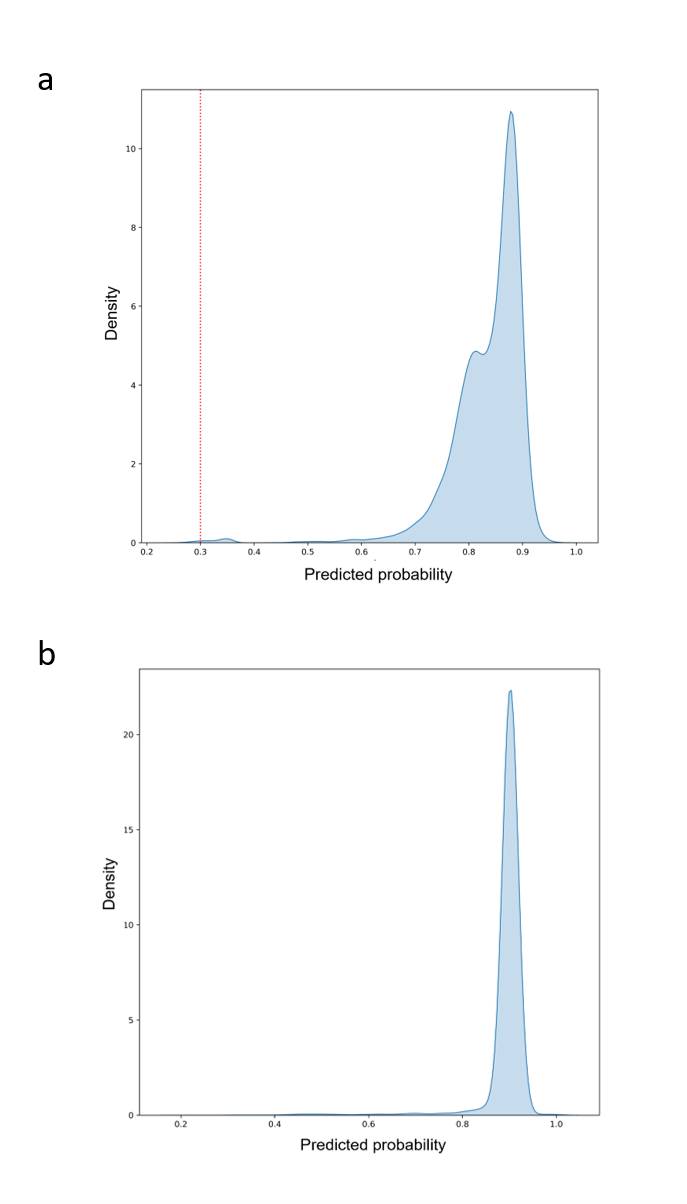

**Supplementary Figure 7. The distribution plot of predicted probabilities.** (a) For the Lee dataset obtained from patients infected with COVID-19. The original test set contains a small number of neutrophils which are not included in the reference dataset. When selecting the 1% cells with the lowest predicted probability, 72 out of the 89 target neutrophils can be filtered out. (b) Distribution plot of a normal PBMC dataset Yazar, which only contains samples from healthy donors.

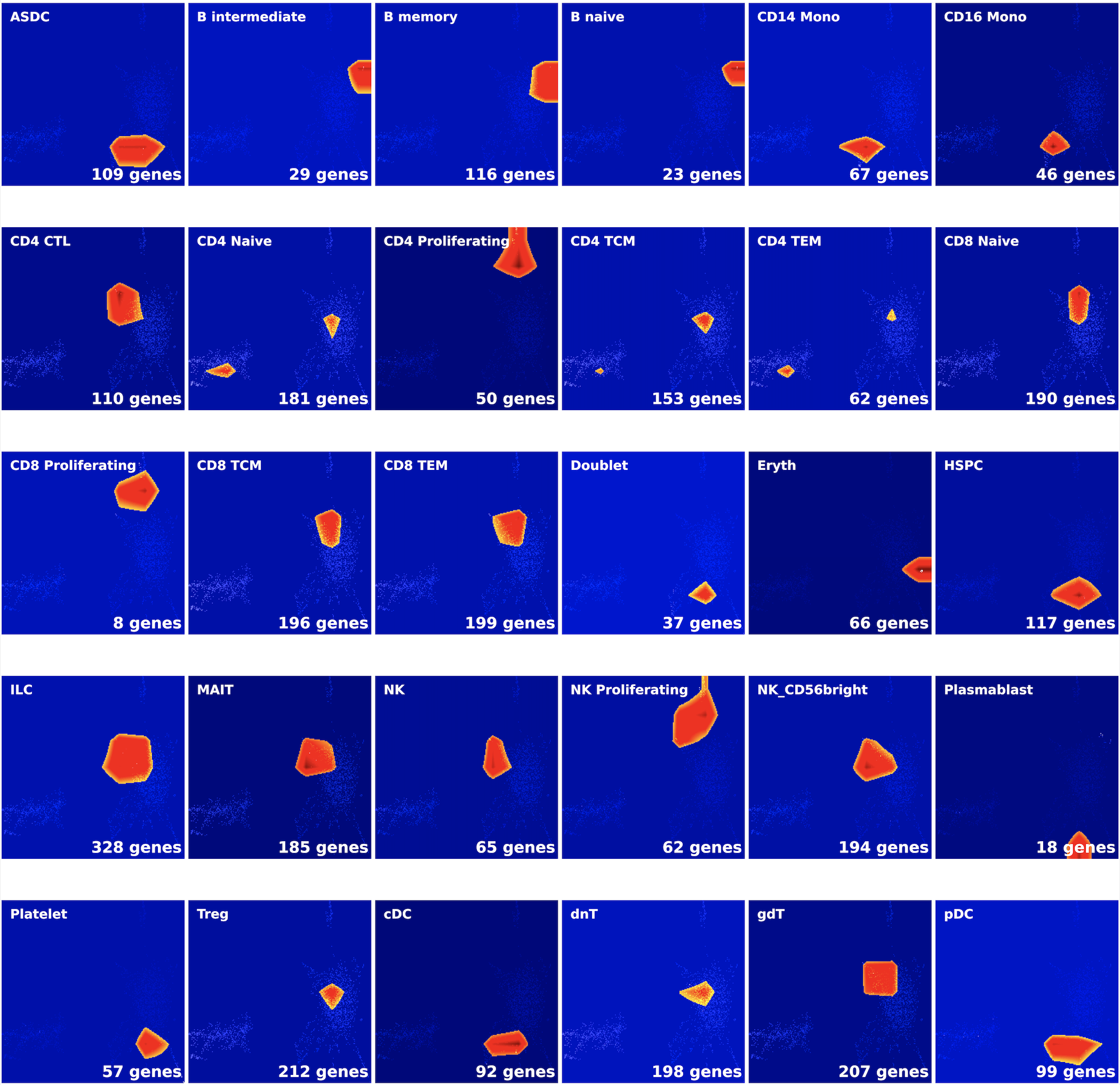

**Supplementary Figure 8. The extracted CAM figures using DeepFeature.** The reference dataset was used for extracting these marker genes. The flatten method is mean and the threshold used for DeepFeature is 0.75. Regions selected using CAM represent highly expressed features in these cell-types. For images generated by DeepInsight, genes have been mapped to pixels, so the extracted features are actively expressed genes in these cell-types. The extracted genes contain multiple marker genes, and the darker the red color, the higher the degree of expression. Compared with the red area, the expression level of the gene in the orange or yellow part is relatively lower.

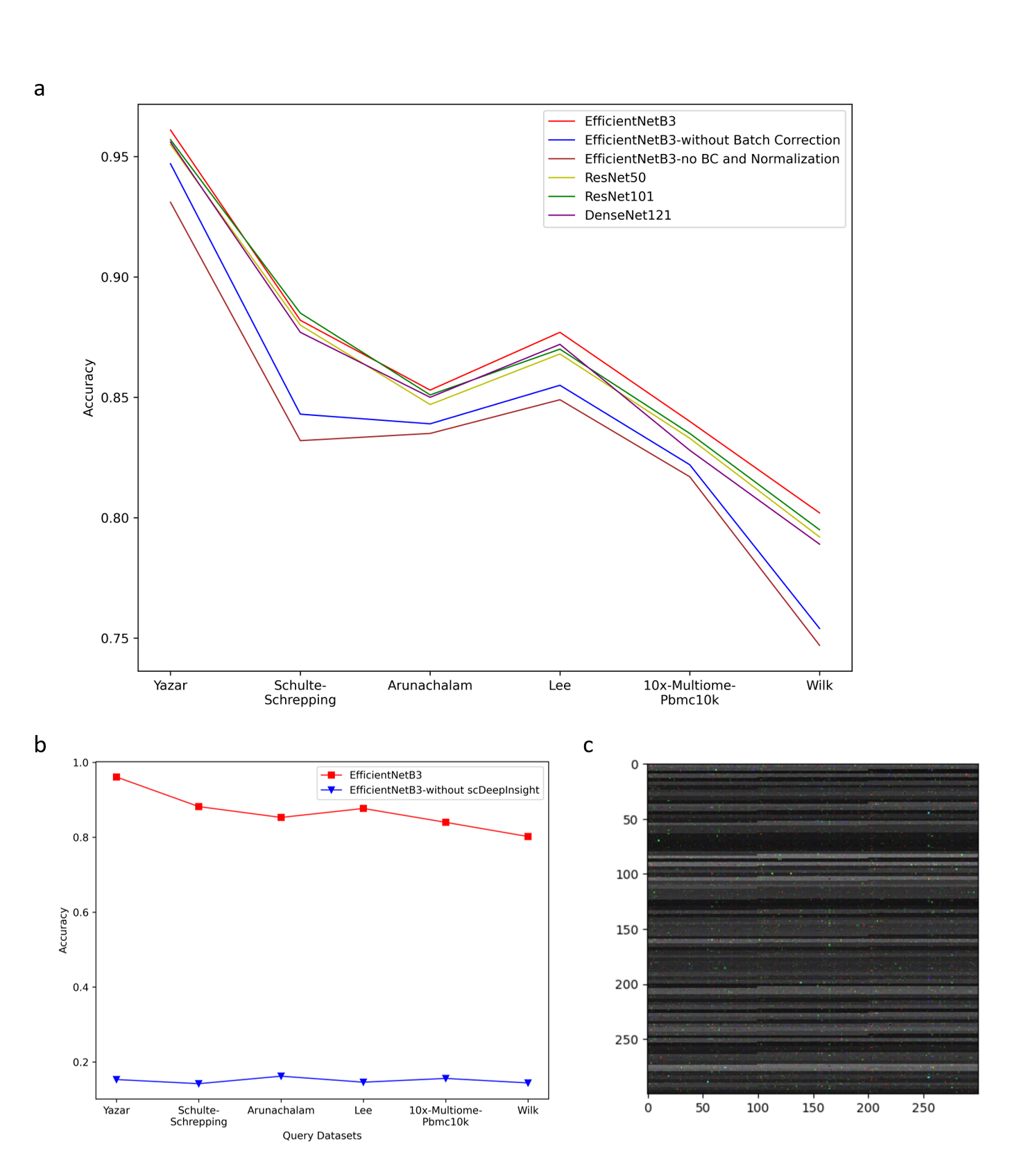

**Supplementary Figure 9 Ablation study with six intra-datasets.** (a) The batch effect correction process improved the accuracy of cell type annotation by an average of 3.18% across six test datasets. When removing the data normalization process, the average accuracy dropped by 0.82%. (b) The average accuracy dropped sharply to 15.2% after replacing DeepInsight with numpy.resize to directly convert the single-cell expression data into images. (c) The converted 3-channel image (300$\times$300) using numpy.resize.

**Parameter values used for scDeepInsight:**

(1) Data preprocessing;

(a) quality control:

Limitations for nFeature_RNA: above 500 and below 5000

Limitations for percent.mt: below 15

(b) Normalization:

method of the function Seurat::SCTransform: "glmGamPoi"

number of variable features: 3000

vars.to.regress: "percent.mt"

(c) Batch effect correction;

normalization.method of the function Seurat::FindIntegrationAnchors: "SCT"

(2) Image conversion process;

perplexity of t-SNE: 50,

distance function (metric) of t-SNE**:** 'euclidean'

learning_rate of t-SNE**:** "auto"

image size of the DeepInsight image transformer: 224*224

(3) Model training process:

The InitialLearningRate: 3e-4.

The weight_decay parameter of L2 regularization: 1e-6

Early stopping epochs: 30

**Parameter values used for the other methods in the benchmarking test:**

(a) SC3:

ks of function sc3: 15

Whether to calculate biology features in sc3: True

(b) FindClusters:

dim of the function Seurat::FindNeighbors: 10

resolution of FindClusters: 0.8

(c) SCINA:

max_iter: 100

convergence_n: 10

convergence_rate: 0.99

sensitivity_cutoff: 1

rm_overlap: 1 (TRUE)

alow_unknown; 1 (TRUE)

(d) Celltypist:

The pretrained model used: Immune_All_Low.pkl

The probability cutoff:0.5

(e) SingleR:

de.method: “wilcox”

(f) scBERT:

The threshold for predicted probabilities: 0.5

num_tokens: 7

dim: 200

heads: 10

depth: 6
